# A unitary model of auditory frequency change perception

**DOI:** 10.1101/2022.06.16.496520

**Authors:** Kai Siedenburg, Jackson Graves, Daniel Pressnitzer

## Abstract

Changes in the frequency content of sounds over time are arguably the most basic form of information about the behavior of sound-emitting objects. In perceptual studies, such changes have mostly been investigated separately, as aspects of either pitch or timbre. Here, we propose a unitary account of “up” and “down” subjective judgments of frequency change, based on a model combining auditory correlates of acoustic cues in a sound-specific and listener-specific manner. To do so, we introduce a generalized version of so-called Shepard tones, allowing symmetric manipulations of spectral information on a fine scale, usually associated to pitch (spectral fine structure, SFS), and on a coarse scale, usually associated timbre (spectral envelope, SE). In a series of behavioral experiments, listeners reported “up” or “down” shifts across pairs of generalized Shepard tones that differed in SFS, in SE, or in both. We observed the classic properties of Shepard tones for either SFS or SE shifts: subjective judgements followed the smallest log-frequency change direction, with cases of ambiguity and circularity. Interestingly, when both SFS and SE changes were applied concurrently (synergistically or antagonistically), we observed a trade-off between cues. Listeners were encouraged to report when they perceived “both” directions of change concurrently, but this rarely happened, suggesting a unitary percept. A computational model could accurately fit the behavioral data by combining different cues reflecting frequency changes after auditory filtering. The model revealed that cue weighting depended on the nature of the sound. When presented with harmonic sounds, listeners put more weight on SFS-related cues, whereas inharmonic sounds led to more weight on SE-related cues. Moreover, these stimulus-based factors were modulated by inter-individual differences, revealing variability across listeners in the detailed recipe for “up” and “down” judgments. We argue that frequency changes are tracked perceptually via the adaptive combination of a diverse set of cues, in a manner that is in fact similar to the derivation of other basic auditory dimensions such as spatial location.

## Introduction

One of the most fundamental tasks of the auditory system is to track changes across sounds as they form sequences over time. For instance, in speech, the “ups” and “downs” of prosody carry information about emotion, about whether a sentence is affirmative or interrogative, or even about the meaning of words in tonal languages [1]. In music, the “ups” and “downs” of melodies constitute the major structural factor for most musical idioms around the world [2]. Both prosody and melody are traditionally described as perceptual “pitch” patterns, isomorphic to changes in the acoustic periodicity of sounds. However, in spite of centuries of intense scrutiny, the precise nature of the auditory cues determining pitch in the general case has remained elusive, with many additional candidates beyond periodicity [3–5]. Moreover, in the perceptual domain, the independence of pitch from other perceptual attributes such as timbre has been recently challenged [6–10]. Here, we explore a unifying hypothesis, suggesting that a salient perceptual dimension on which sounds can be ordered from “low” to “high” is in fact a compound construct, seamlessly derived by listeners from the combination of distinct auditory cues, each adaptively weighted in a manner that can be sound-dependent and listener-specific.

In the following experiments and modeling, we investigated two main types of acoustic cues: spectral fine-structure cues (SFS) and spectral envelope cues (SE). Such a dichotomy is in part motivated by the standard source-filter model of sound production [11, 12], whereby an excitatory source generates a signal which is then filtered by resonant bodies. For the voice, the source would be vocal cords and the filter the vocal tract. For a musical instrument such as the violin, the source would be strings and the filter the body of the violin. There is an approximate mapping between source and SFS on the one hand, and filter and SE on the other hand. Sources constrain SFS, for instance for periodical excitatory motion that produce harmonically-related frequency components with frequencies at multiples of a fundamental frequency (*f*_0_). In contrast, filters constrain SE, through resonances that shape the global amplitude profile of the frequency spectrum. Our choice of SFS and SE is also motivated by perceptual considerations. Harmonicity in SFS, equivalent to periodicity, has long been associated to pitch, and may also explain the pitch of inharmonic sounds [13]. SE cues are thought to affect timbre and source recognition [14, 15]. In particular, the balance of low vs. high energy imposed by SE is associated to a timbre dimension termed brightness [16, 17]. However, as we will now briefly review, there is also a growing amount of evidence that SFS and SE interact for pitch perception, and that the distinction between pitch and brightness may not be as clear-cut as initially thought.

The most basic definition of pitch is “that attribute according to which sounds may be ordered on a scale extending from low to high” [18, 19]. The idea that such a scale may be constructed from more than one single acoustic cue is not new. For harmonic sounds, Bachem [20] introduced the now classic distinction between “pitch chroma”, a circular dimension repeating at the octave, and “pitch height”, a linear dimension following absolute frequency (see also [13, 21]). Pitch has been further qualified, with notions of “virtual pitch”, “spectral pitch”, “melodic pitch” introduced for various combinations of SFS and SE cues [22–24]. For inharmonic sounds, recent results by McPherson and McDermott add yet additional candidate cues to try and explain pitch judgements [25]. Using a variety of tasks usually understood as involving pitch, such as pitch discrimination, melodic contour identification, or interval judgments, the authors concluded that pitch judgements were in fact based on multiple mechanisms operating concurrently, such as the computation of *f*_0_ from SFS cues for harmonic sounds, and the tracking of SE cues for inharmonic and harmonic sounds (see also [26]). Previously, frequency-shift detectors tracking local changes in SFS cues had also been evidenced [27], providing yet another mechanism contributing to “up” or “down” pitch judgements available for harmonic and inharmonic sounds alike [28].

Another line of evidence suggesting that pitch judgments may be based on more than one type of acoustic cue is found in studies where SFS and SE cues were combined in an antagonistic manner. The experimental findings using such paradigms appear contradictory so far. Small simultaneous SFS and SE changes can be processed largely independently by some listeners at least [29, 30] while timbre judgments involving SE are largely unaffected by changes in SFS [31]. However, it is also the case that SFS and SE cues interfere in discrimination experiments with adult listeners, such that simultaneous variation in timbral brightness impairs pitch discrimination, and vice versa [8, 32–35]. Interestingly, there exist recent reports of 3 and 7 month infants being “immune” to the interaction of SFS and SE [36], suggesting a role for long-term auditory learning in the interaction [37, 38]. One ecological interpretation for such a learnt interaction between SFS and SE may be the actual coupling of *f*_0_ and SE observed for natural sounds, underlining limitations of the classical source-filter physical model [39].

Finally, an intriguing synergy between SFS and SE cues has been revealed through the crossmodal correspondence of auditory pitch with the spatial dimension of height. Empirically, it has been shown that “high” pitches are spontaneously associated with high spatial elevation [40, 41]. However, while the association is strong in Western musicians and parallels the way Western musical notation represents pitch, recent work suggests that listeners without musical training only exhibit such spatial association effects for frequency changes consisting of coupled SFS and SE shifts, but not for one-dimensional shifts alone [6].

We now turn to the distinction that is conventionally made between pitch and timbral brightness. An intuitive definition of pitch, in addition to the “low” to “high” scale, is that it is “that attribute of sounds of which melodies are made of” [23, 42]. However, pitch may not be the only perceptual dimension with which melodies can be made of. A number of studies have now revealed strong and unexpected commonalities between the perception of SFS and SE in sequences akin to melodies. These include the recognition of familiar tunes based on brightness alone [43] or the accurate discrimination of melodic patterns for both pitch and brightness [28, 44, 45]. These commonalities stand in sharp contrast with the qualitative differences observed between e.g. pitch and loudness sequences [46, 47]. Furthermore, even though melodies are only conceived in terms of pitch changes for Western tonal music, musical idioms outside of Western culture do use “melodies” made out of timbral changes, such as throat singing or jaw’s harp in East Asia, or the musical bow in Africa [48]. In all of these musical techniques, a fixed SFS is being modulated by changes in SE, incompatible with the traditional Western musical definition of melody. These observations cast further doubt on the long-held but perhaps arbitrary distinction made between pitch and brightness as separate perceptual correlates of sound sequences with dynamically-changing frequency content.

In the present study, we introduce a generalized version of Shepard tones [49] to further address the role of SFS and SE cues on judgements of frequency change on a “low” to “high” scale. Shepard tones were originally designed to tear apart the contributions of chroma and pitch height cues to pitch perception [21, 49–51]. They consist of octave-spaced pure tone components, defining chroma, with a fixed bell-shaped spectral envelope, defining height. A sequence of Shepard tones with increasing chroma is cyclic, as a full octave shift is acoustically identical to no shift at all. Such sequences give rise to the famous Shepard scale illusion, that is, a sequence of tones with seemingly continually ascending or descending pitch. Shepard tones were also found to produce ambiguous frequency shift judgements for half-octave shifts. Typically, listeners’ responses were split between upward and downward shifts for the same sound in this so-called tritone paradox [49, 51]. Finally, the pitch percept associated to ambiguous shifts between Shepard tones can be strongly biased by preceding context [52, 53]. In the original Shepard tones, only SFS is manipulated, while SE is kept constant. Interestingly, exactly analogous findings for all of these observations were recently reported using SE changes instead of SFS changes. Sounds were designed to have local SE shifts combined with a fixed, harmonic SFS. This produced a Shepard scale illusion, a tritone paradox, and context effects for SE cues [45].

The further generalization of Shepard tones we introduce here aims at independently manipulating SFS and SE cues, in a symmetric and cyclic manner, both for harmonic and inharmonic sounds (Fig. 1). We used the trick of even-to-odd harmonic attenuation [54, 55] to obtain a circular SFS dimension with harmonic sounds. These tones were coupled with an SE that was quasi-periodic along the frequency axis [45]. Combining the resulting octave-circular SFS and octave-circular SE dimensions meant that the two dimensions were intrinsically comparable with regards to the size of shifts along each dimension. By jittering the frequency position of partial tones [25], stimuli could also be made inharmonic.

**Fig 1.**
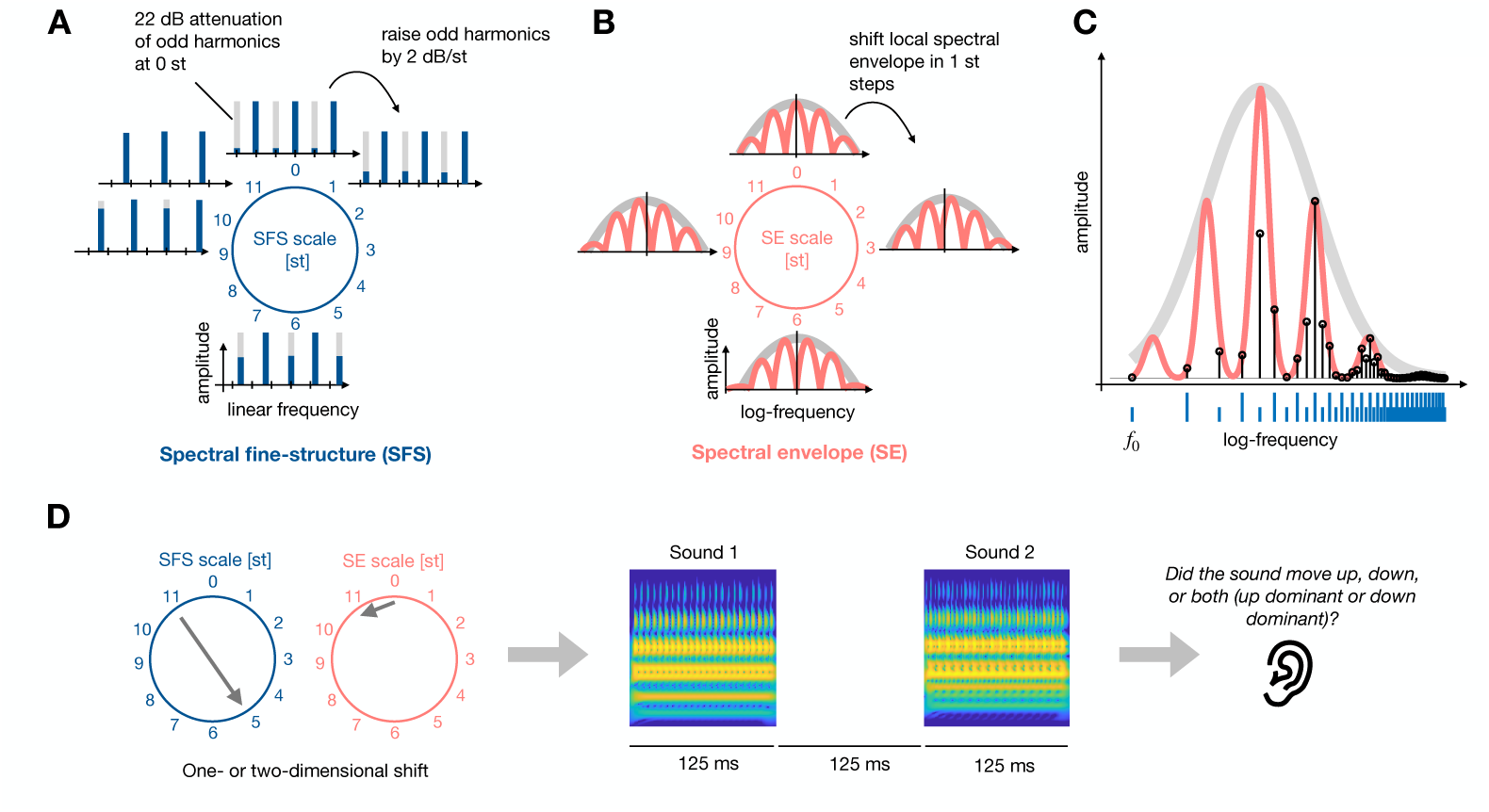
Stimulus schematic for the generalized Shepard tones used in Exp. 1 and 2. (A) Spectral fine structure (SFS, blue) is illustrated here as a set of harmonic components at integer multiples of a fundamental frequency *f*_0_. SFS could be shifted in 1-semitone (st) steps, while manipulating the amplitude of odd harmonics at each step to induce perceptual circularity across a 12-ST shift. Inharmonic stimuli were created by jittering the individual frequency components around their nominal values. (B) Spectral envelope (SE, red) had peaks at octave-related frequencies superimposed with a fixed bell-shaped envelope. SE could also be shifted in st steps and displayed exact acoustic circularity across a 12-ST shift. (C) Exemplary stimulus for a given combination of SFS and SE. Frequency components are constrained by the SE (red line) and follow the harmonic series of the SFS and its even-odd attenuation (indicated by blue lines). The resulting amplitude of components is shown by black lines. Note that some components are lower than their nominal SE amplitude, because of the even/odd attenuation. (D) Schematic depiction of exemplary trial with two-dimensional shift of 6 st in the SFS dimension and 11 st in the SE dimension. After each trial, participants were asked to to judge whether the sound pair moved up, down, both (up dominant), or both (down dominant). Stimuli for Exp. 3 were generated in a similar manner except that the SFS was not harmonic, but with fixed distance on a log-frequency scale, allowing for exact acoustic circularity for a shift corresponding to their log-frequency distance.

Experiment 1 tested whether these generalized Shepard tones yielded classic Shepard properties. Participants were asked to report “up” or “down” shifts across pairs of tones. Such subjective changes were described to participants as either changes in pitch or brightness, to test our core hypothesis that, irrespective of vocabulary, consistent “up” or “down” judgments could be made irrespective of the cue manipulated. Experiment 2 tested interference between SFS and SE shifts presented concurrently, in a synergistic or antagonistic manner, for harmonic and inharmonic tones. Importantly, to test whether two explicit perceptual dimensions were simply competing during the judgements, participants had the opportunity to report whether they perceived “both” directions of change concurrently. This rarely happened. Experiment 3 tested whether the observed effects generalized to an SFS with a different spacing of partial tones. Finally, to provide a synthetic interpretation of the rich but complex behavioral dataset and quantify the auditory cues compatible with the observed perceptual data, we used a computational model combining both SFS-related and SE-related cues. Fitting the model to each listener across all experiments revealed common stimulus-based factors as well as inter-individual effects, characterizing the recipe for “up” vs. “down” judgments in a novel degree of detail and suggesting that auditory frequency change perception is based on an adaptive combination of a diverse set of cues.

## Results

### One-dimensional shifts of harmonic and inharmonic sounds

In Exp. 1, we tested the perception of one-dimensional SFS or SE shifts for generalized Shepard tones as described in Fig. 1. Pairs of sounds to be compared perceptually were presented on each trial, with shifts of SFS or SE between 1 and 11 semitones (st) introduced across the two sounds of the pair. Because of the design of the stimuli, a shift of 12 st is equivalent to no shift at all, so all possible shift values were effectively sampled at a 1 st resolution. In addition, harmonic and inharmonic SFS were both tested. Inharmonic SFS was obtained by jittering the nominal frequency values of individual frequency component of the harmonic SFS case, keeping the jitter values fixed across the two sounds within a trial. Blocks of trials were constructed for each stimulus type (SFS harmonic, SFS inharmonic, SE harmonic, SE inharmonic) and all shift values were randomly presented within a block. Listeners were asked to report whether they heard the sound pair moving “up” or “down”, with the instructions specifying that the changes could be characterized as either pitch or brightness.

Results are shown in Fig. 2. As expected, we obtained a general trend of increasing “down” responses as a function of shift size in all cases. In most cases, the increase was monotonic, with the most notable exception being the inharmonic SFS shifts. In the latter condition, we observed an increase of the proportion of “down” responses up to 6 st, followed by an unexpected u-shaped pattern of scores from 6 to 11 st. For SE shifts, there also was a tendency to plateau or even reverse the general trend for extreme shifts smaller than 2 st or larger than 10 st. In all cases, however, the results displayed a hallmark feature of Shepard tones: shifts of 6 st yielded ambiguous responses (around 50% “down”, 50% “up”), thus providing a first indication that our generalization of Shepard tones to SFS and SE shifts, harmonic or inharmonic, was successful.

**Fig 2.**
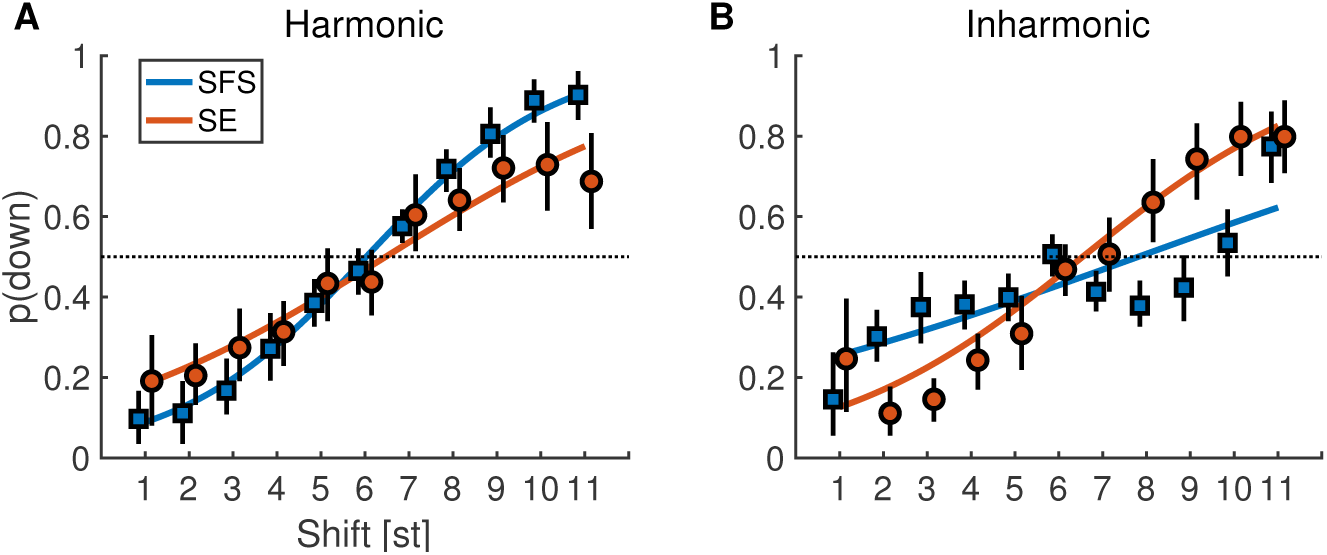
Proportion “down” responses for one-dimensional shifts of Exp. 1 for harmonic (A) and inharmonic (B) Shepard sounds. Colored lines correspond to the fitted GLME model. Black vertical lines indicate 95% confidence intervals obtained by bootstrapping.

The raw data were fitted by a general linear mixed-effects (GLME) model (see Methods) . The colored lines in Fig. 2 correspond to the fixed-effects predictions of the GLME model. Specifically, the model did not indicate a main effect of shift type, SFS vs. SE (*β* = 0.01 [−0.08, 0.1], *p* = .87) but confirmed a clear effect of shift size (*β* = 0.31 [0.30, 0.32], *p* < .001). Further, the GLME confirmed interactions between harmonicity and shift type (*β* = −0.52 [−0.61, −0.43], *p* < .001), harmonicity and shift size (*β* = 0.06 [0.04, 0.07], *p* < .001), as well as harmonicity, type, and size (*β* = 0.1 [0.08, 0.11], *p* < .001). The latter interaction corresponded to the shallower slopes for SE shifts compared to SFS shifts for harmonic sounds and the reversal of that pattern for inharmonic sounds. This three-way interaction indicated that harmonicity mediated the strength of the shift percept for one-dimensional SFS and SE shifts: SFS shifts were more clearly discernible for harmonic sounds whereas SE shifts became clearer for inharmonic sounds.

### Two-dimensional shifts of harmonic and inharmonic sounds

In Exp. 2, we set out to explore the perception of two-dimensional shifts with SFS and SE combined, and sometimes pitted against each other. In this experiment, shifts were concurrently introduced in both SFS and SE, with independent shift values for SFS and SE. We sampled three shift values for each type of cue: 1, 6, and 11 st, providing nine different 2D combinations of shifts for either harmonic SFS or inharmonic SFS.

Importantly, on each trial, listeners were allowed to report whether they perceived both directions of change concurrently. Four response possibility were thus provided: “up”, “both (up dominant)”, “both (down dominant)”, and “down”. If listeners explicitly weighed pitch and brightness judgements to provide their response, then a large proportion of “both” responses should be observed when SFS and SE shifts were in opposite directions. Conversely, if they simply used a perceptual “low” to “high” scale by implicitly combining SFS and SE cues on a unitary perceptual dimension, little or no “both” responses should be observed. Results showed that “both” responses were only observed for a relatively small fraction of the trials (on average 8 and 14 percent of overall responses for synergistic and antagonistic trials with harmonic sounds, respectively; and 10 and 19 percent of responses for synergistic and antagonistic trials with inharmonic sounds; see Supplementary Fig. 1). Responses were thus aggregated as either “up” (“up” together with “both (up dominant)”) or “down” (“down” together with “both (down dominant)”) for comparison with Exp. 1. The resulting patterns of response are shown in Fig. 3. Visual inspection of this Fig. 3 suggests that the effect of SE shifts was smaller than the effect of SFS shifts for harmonic sounds, but SFS and SE shifts had similar effects for inharmonic sounds. Interestingly, shifts of 6 st along the SE dimension (condition “SE6”) appeared to yield a reduced effect of SFS for harmonic sounds and a complete lack of an effect for inharmonic sounds (Fig. 3B). Furthermore, incongruent SFS and SE shifts appeared to exactly cancel out for inharmonic sounds, with SFS1-SE11 and SFS11-SE1 shifts producing around 50% “down” responses.

**Fig 3.**
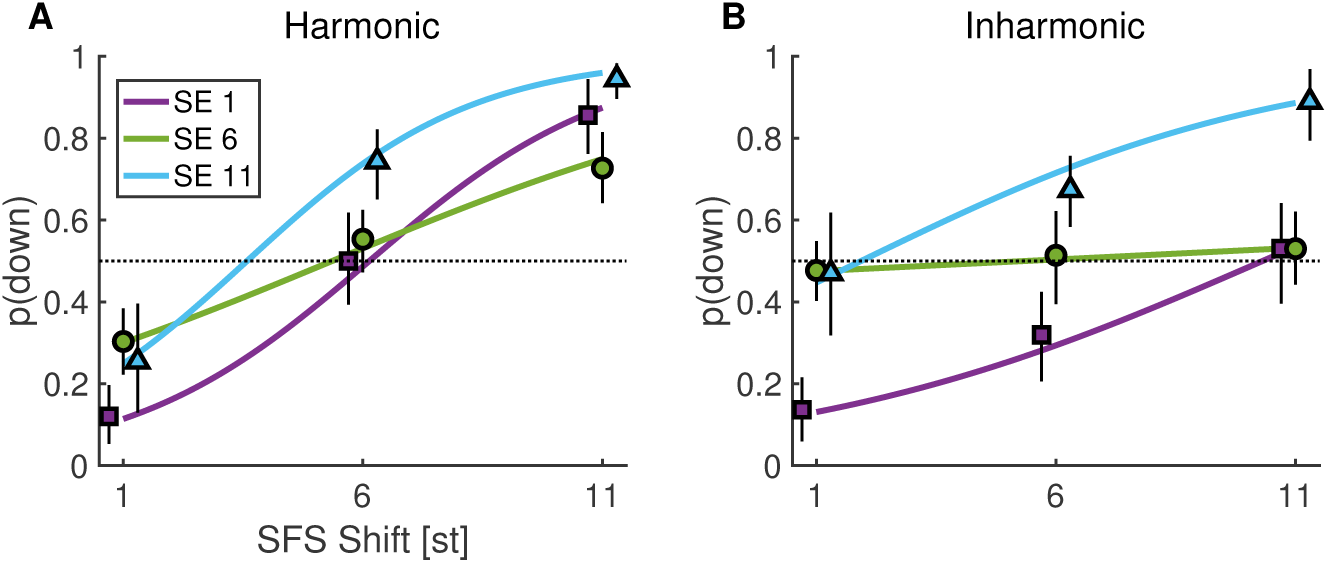
Proportion “down” responses for two-dimensional shifts of Exp. 2 for harmonic (A) and inharmonic (B) Shepard sounds. Colored lines indicate the fixed effects of the GLME model. Black vertical lines indicate 95% confidence intervals obtained by bootstrapping.

A similar GLME model as in Exp. 1 was fitted to the data. The model suggested that listeners tended to perceive harmonic sound pairs more frequently as moving “down” compared to inharmonic shifts (*β* = −0.39 [−0.49, −0.3], *p* < .001). There were strong main effects of SFS (*β* = 0.25, [0.23, 0.26], *p* < .001) and SE (*β* > 0.7, [0.570.83], *p* < .001), the latter parametrized by the two factors SE1/11 and SE6/11 that used effects coding to contrast conditions SE1 and SE6 with SE11, respectively. Different types of regressors were used for SFS and SE variables in order to account for the interactions of SFS and SE factors with different slopes of SFS for the SE6 level compared to the other SE levels (see Fig. 3 and next paragraph for an alternative model). Importantly, the effect of SFS shifts were markedly greater for harmonic sounds compared to inharmonic sounds as indicated by a harmonicity-SFS interaction (*β* = 0.09 [0.08, 0.11], *p* < .001). To the contrary, the difference between the SE1 and SE11 conditions were enhanced for inharmonic sounds as indicated by a harmonicity-SE1/11 interaction (*β* = 0.22 [0.07, 0.37], *p* = .004). Further, the SFS factor interacted with both the SE1/11 factor (*β* = 0.06 [0.03, 0.08], *p* < .001) and the SE6/11 factor (*β* = −0.14 [−0.16, −0.12], *p* < .001), which reflected the drastically reduced effect of SFS in the SE6 condition, that was even close to zero for inharmonic sounds.

The GLME model above used different predictors for the SFS and SE factors to account for interactions. To quantitatively compare both factors, we also ran a GLME with numerical predictors for both variables. This model indicated that SFS shifts indeed yielded larger main effects (*β* = 0.22 [0.20, 0.25], *p* < .001) compared to SE (*β* = 0.12 [0.10, 0.14], *p* < .001). At the same time, SFS shifts also depended positively on harmonicity (*β* = 0.09 [0.07, 0.12], *p* < .001), whereas the harmonicity-SE interaction was weaker and in the opposite direction (*β* = 0.03 [−0.06, −0.01], *p* = .005). Thus, SFS had stronger effects compared to SE, but the effect of SFS was reduced for inharmonic sounds, whereas the effect of SE was only slightly greater for inharmonic sounds compared to harmonic sounds.

In sum, the two-dimensional shifts of Exp. 2 showed a dominance of SFS cues for harmonic sounds, but an increase of the contribution of SE cues for inharmonic sounds. This is coherent with the one-dimensional shifts of Exp. 1, where SFS responses had steeper slopes than SE responses for harmonic sounds, whereas SE responses had steeper slopes than SFS responses for inharmonic sounds. Importantly, for inharmonic sounds, we observed a cancellation effect of incongruent SFS and SE shifts. To confirm that this pattern of responses was not due to the specific type of inharmonicity induced by the linear jittering operation together with the odd-even attenuation, we conducted Exp. 3 using a potentially more simple, yet still inharmonic partial series.

### One- and two-dimensional shifts with log-equidistant partials

In Exp. 3, we replaced the linearly-spaced partial series by a partial series, in which all partials were spaced 4 st apart (i.e., equidistant on a logarithmic frequency scale), such that there were three partials per octave and the resulting SFS shift was 4-st periodic. This can be viewed as a compressed version of the octave-spaced original Shepard tones. Such a choice was motivated by the need to sample accurately-enough the SE cues over a reasonable range of audible frequencies. As a consequence, only three different SFS shifts could be tested: 1 st (up), 2 st (ambiguous), and 3 st (down). These are conceptually equivalent to the three shifts tested in Exp. 2. The SE cues were kept identical as for Exp. 1 and 2, with an octave spacing between peaks and hence 12-st circularity. For the SE cues, shifts of 1 st (up), 6 st (ambiguous), 11 st (down) were used. Exp. 3A paralleled Exp. 1, as we tested one-dimensional shifts with 1, 2, 3 st shifts for the SFS cues or, separately, 1, 6, 11 st shifts for the SE cues. Exp. 3B paralleled Exp. 2, as we tested 2D shifts along both the SFS and SE cues concurrently. The methodological details were otherwise identical between Exp. 1 and Exp. 3A, and between Exp. 2 and Exp 3B.

Figure 4 shows the results for both Exp. 3A and 3B, following the convention of previous figures. For Exp. 3A, the GLME fit indicated an effect of shift type, SFS or SE (*β* = −0.92 [−1.24, −0.61], *p* < .001) and shift size in st (*β* = 1.26 [1.11, 1.41], *p* < .001). Both effects are qualified by a strong interaction between type and size (*β* = 0.52 [0.37, 0.67], *p* < .001), suggesting a stronger effect of SFS cues compared to SE cues. For Exp. 3B, again we observed a relatively small fraction of trials for which “both” responses were provided (on average 13 percent of responses for synergistic trials and 23 percent of responses for antagonistic trials, see Supplemental Fig. 1 for details). Using a similar analysis method as for Exp. 2, we again obtained an interaction between the SFS and SE factors. Besides effects of SFS and SE (see Tab. 4 in the Supplementary Materials for details), there was a robust interaction between the SFS and SE6 factor (*β* = −1.0 [−1.14, − 0.86], *p* < .001), corresponding to the lack of effect of SFS shifts in the SE6 condition, but enhanced differences for the SE11 condition. Overall, Exp. 3 confirmed the interaction between SFS and SE cues in perceptual judgments and suggested that SE cues were robust in 2D shifts for a broader class of inharmonic sounds compared to those tested in Exp. 1 and 2.

**Fig 4.**
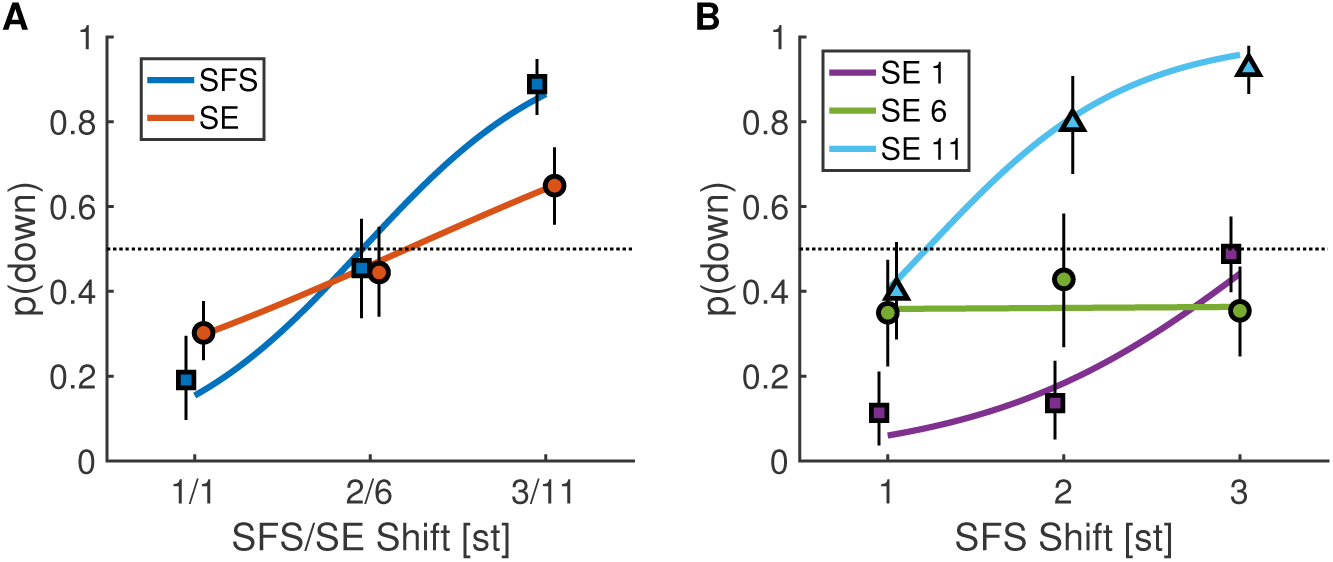
Proportion “down” responses in Exp. 3 with log-equidistant partials for one-dimensional shifts (A) and two-dimensional shifts (B). Colored lines indicate the fixed effects of the GLME model. Black vertical lines indicate 95% confidence intervals obtained by bootstrapping.

## Computational modeling

### Rationale

Our stimuli were constructed along two acoustic dimensions, SFS and SE, and included harmonic sounds and two types of inharmonic sounds. The experimental conditions provided a rich dataset where all of these factors were studied, in isolation and in combination, with interactions between them. To provide a synthetic understanding of the results, we now turn to computational modeling. Based on previous results [5], one may broadly differentiate three types of candidate *auditory* cues (in contrast to the *acoustic* dimensions used for the stimulus design) that could underlie the perceptual judgements of listeners. First, listeners could have extracted a single pitch estimate for each sound in a trial, and compared the estimates across the two sounds of a trial. Such a pitch estimate is broadly assumed to be based on periodicity extraction, or, equivalently, *f*_0_ estimation by matching harmonic templates to the SFS [5]. Second, listeners may have extracted frequency-shifts between individually resolved frequency components, that is, frequency components that are represented by distinct neural populations after peripheral auditory filtering in the cochlea [27, 56]. These frequency-shifts could be computed across pairs of components over the two sounds in a trial by a population of frequency-shift detectors and the dominant output from the population could then be taken as the direction of perceptual shift. Equivalently, such cues from resolved frequency components may also related to so-called spectral tracking [25]. Third, listeners may compare the broad spectral peaks defined by the unresolved frequency components, that is, frequency components exciting overlapping neural populations after peripheral auditory filtering. These unresolved harmonics can provide coarse-grained spectral cues, such as formants, and have been shown to act as an important basis for timbre processing [12, 57, 58].

We implemented a relatively simple computational model to estimate these three types of cues (see Fig. 5 and Methods section for details). For a fair comparison, all cues were estimated after a common front-end simulating peripheral frequency filtering at the level of the cochlea, to provide an “auditory representation” capturing some features of the neural representation available to the auditory system. The front-end chosen was a standard Gammatone auditory filterbank decomposition [59, 60]. The first cue was based on the extraction of peaks in the autocorrelation (AC) function over time at the output of each Gammatone filter, summed across all filters, implementing a standard model of pitch estimation [5]. For the second and third cues, we computed the “excitation pattern” produced by each sound. Excitation patterns are a spectral representation of sound with frequency resolution matched to auditory peripheral filtering. They were computed here as the root-mean square of the output of each Gammatone filter over time. To determine the dominant direction of frequency shift in a given trial, we cross-correlated the excitations patterns for the two sounds of the trial and computed the center of mass (on the frequency axis) of this function. For the second cue, which was broadly conceived to capture resolved partials (CCres), the cross-correlation was computed for the low-frequency part of the excitation patterns. For the third cue, which was conceived to capture unresolved components (CCunres), the cross-correlation was computed for the high-frequency part of the excitation patterns. The frequency limit was parsimoniously chosen at ten times the geometric mean of the range of *f*_0_ used in the experiment (905 Hz), corresponding to the average frequency of the tenth partial component of the test sounds. Other choices of frequency limits yielded qualitatively and quantitatively very similar results, indicating that the model did not critically rely on this choice.

**Fig 5.**
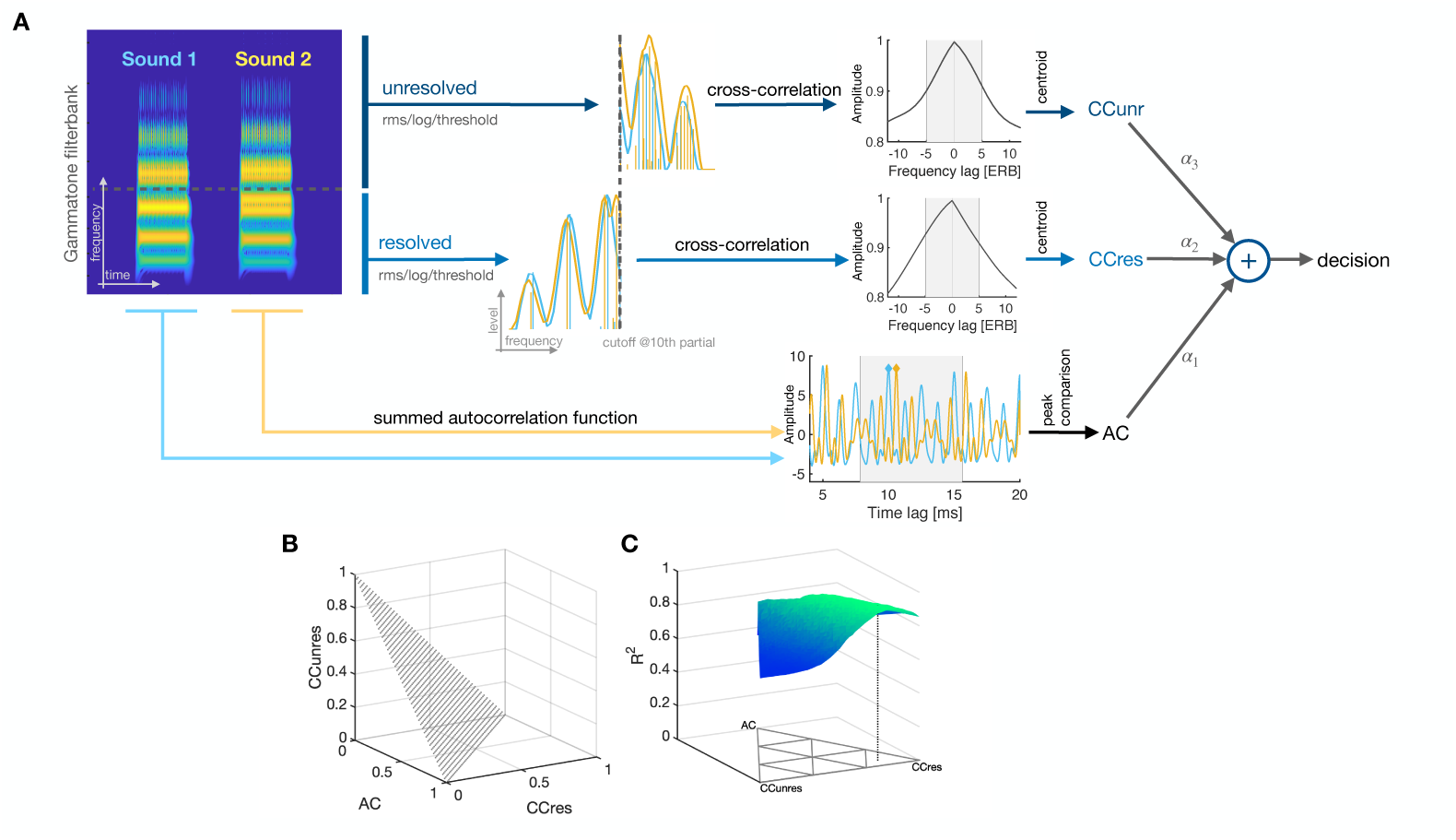
(A) Schematic of cue computation and integration, illustrated with a pair of harmonic sounds shifted by 11 and 1 st along the SFS and SE dimension, respectively. For the AC cue, autocorrelations within each auditory band are computed and subsequently summed across bands. From the resulting summed autocorrelation function, peaks are extracted from a search range corresponding to 64-128 Hz (gray patch) and subtracted, yielding an estimate of the periodicity difference of the two sounds; positive differences indicate downward movement. CC cues are computed using the rms-energy of bands across time, followed by a logarithm (conversion to level-domain) and thresholding below −30 dB peak level. The CCres cue considers the presumably resolved part of the excitation pattern below 905 Hz (geometric mean of the frequency of the 10th partial component across Exps. 1 and 2); the CCunres cue considers the unresolved part above that frequency limit. For both types of cues, the remaining excitation patterns are cross-correlated and the centroid (first moment) is computed in a search range (gray patches). Positive centroids indicate downward movement. Cues are integrated in a weighted sum (positive sum coded as “down”). (B) Possible cue weightings of sum one. (C) Cue weights projected onto x-y plane; the farther a point’s distance to a specific edge, the less weight the respective cue receives. The corresponding model fit is displayed on z-axis for one human listener’s results in the inharmonic SFS-EN condition. The dashed line shows the optimal weights for this example.

These three types of auditory cues, AC, CCres, and CCunres, can be approximately mapped to the SFS and SE acoustic dimensions manipulated in the experiments. The AC cue should be impacted by changes along both SFS and SE dimensions. The CCres cue focuses on a region that is known for having a dominant impact on pitch [5]. Cross-correlation is not a standard way to estimate pitch, however, so our computation may be best understood in terms of local frequency-shift detectors based on SFS, rather than as a global pitch estimate. The CCunres cue is the one most directly related to SE as conceived in timbre models of e.g. brightness [16]. To check for these a priori mappings, correlations between the three different auditory cues, and between the cues and behavioral data, will be presented together with the modeling results.

Finally, because the behavioral data showed that listeners mostly provided a unitary estimate of “up” or “down” shift, we integrated all three cues in a decision model. To this aim, the three cues were normalized by their variance and integrated as part of a weighted sum, such that the model yielded a single “up” or “down” decision on every trial. The model was then fitted by optimizing the correlation between the model and individual participants per experimental condition. The optimization procedure was a simple grid search. Fig. 5B depicts the resulting parameter space and Fig. 5C illustrates the grid search.

### Modeling results

Across all experiments, all three auditory cues turned out to be relatively independent from each other. Computing the inter-cue correlation across all simulated trials of the three experiments indicated no relation between the AC cue with any of the two CC cues (*p* > .26). A weak, yet non-negligible linear relation was only observed between CCres and CCunres (*r* = .22, *p* < .0001). The low inter-cue correlation thus validates the a priori hypothesis that AC, CCres, CCunres cues captured qualitatively different aspects of the experimental stimuli.

The accuracy with which individual behavioral response patterns could be explained by a combination of the three auditory cues was generally good, although it varied as a function of the stimulus condition, as seen in Fig. 6A-C. For harmonic SFS shifts, model correlations with individual data turned out to be higher compared to inharmonic SFS shifts. Conversely, for harmonic SE shifts, model accuracies were lower compared to inharmonic SE shifts. For two-dimensional SFS-SE shifts, the fitted models generally closely matched individual response patterns. We did not quantitatively evaluate Exp. 3A due to the small number of shift conditions. To illustrate which aspects of the behavioral response patterns were captured by the model, Fig. 7 presents the average response patterns from listeners together with the model average across all individually fitted models. The harmonic conditions, for SFS, SE, or two-dimensional SFS-SE shifts, yielded excellent fits with beyond 95% of shared variance between model and human data. In the inharmonic conditions, we still obtained more than 86% of shared variance for SFS shifts and more than 90% shared variance for SE and SFS-SE shifts. The most challenging condition to interpret with the model appeared to be Exp. 3B, with around 75% of shared variance. Furthermore, all three types of cues contributed in various combinations across experimental conditions and listeners. This is demonstrated in Supplementary Fig. 2, depicting paired differences of model fit between the full model and reduced model variants with fewer cues. The chosen model dimensionality thus seems necessary to fully account for the present data.

**Fig 6.**
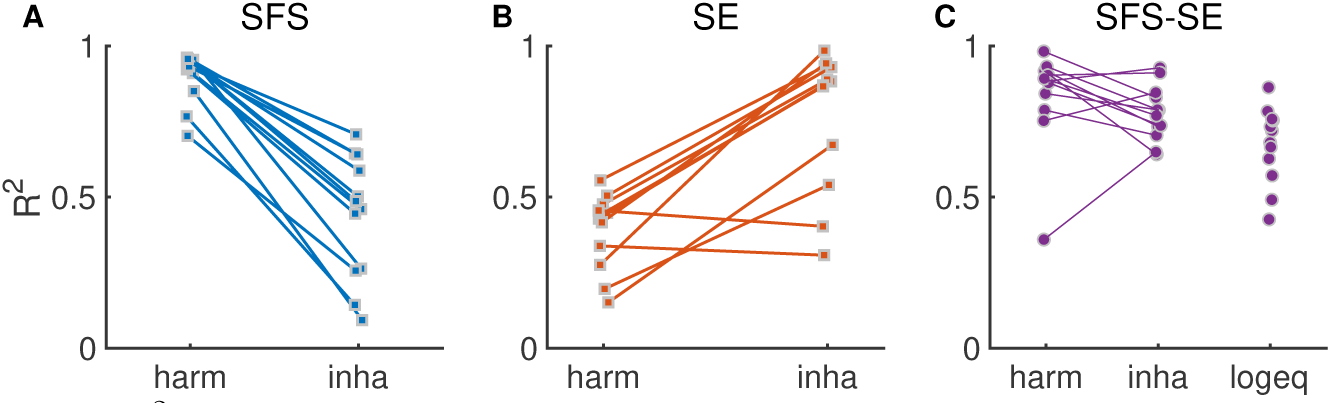
(A) *R*^2^ values of weighted model and empirical data of individual participants for SFS shifts of Exp. 1 with harmonic (harm) and inharmonic (inha) partial series. (B) Same as (A) but for SE-shifts. (C) Same as (A-B) but for the two-dimensional SFS-EN shifts of Exp. 2 (harm & inharm) and Exp. 3 (log-equidistant-spaced partial series).

**Fig 7.**
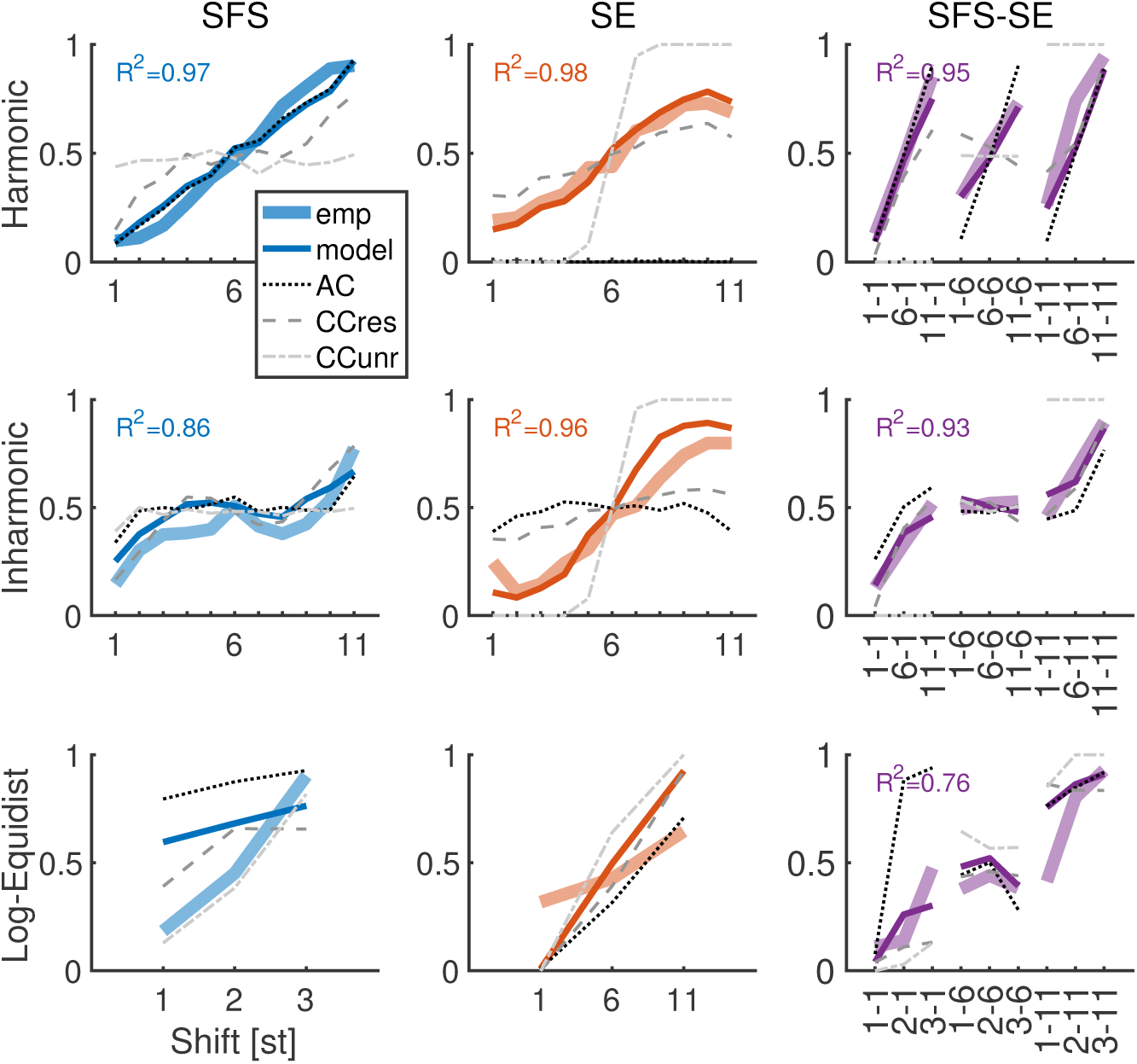
Model results and human listeners across all experiments. The y-axis shows the proportion of “down” responses. The x-axis indicate shift size; for two-dimensional shifts, pairs indicate shifts along the SFS and SE dimension, respectively. The rows index the type of fine structure (harmonic, inharmonic (Exps. 1&2) and log-equidistant (Exp. 3). The columns index the type of shift, one-dimensional SFS or SE, and two-dimensional SFS-SE with average data from human listeners and the model plotted in thick and thin colored lines, respectively. Raw predictions from the AC, CCres, and CCunres cues as indicated in the legend. *R*^2^ values for correlation between average model and empirical data (omitted for log-equidistant SFS and SE shifts of Exp. 3 due to insufficient number of shifts). *R*^2^ values for raw cues are provided in Tab. 5 in the Supplementary Materials.

Our modeling approach allowed us to investigate which cues enabled such high shared variance between the behavioral and modeling data, for each experimental condition. The corresponding strength of the correlations between average human data and raw auditory cue estimates is provided in Supplementary Tab. 5. Overall, the CCres cue appeared to be most versatile, yielding significant correlations for all of the three shift types, SFS, SE, and SFS-SE. The CCunres cue only yielded significant correlations for one-dimensional SE shifts. The AC cue yielded high correlations for SFS and SFS-SE shifts but not for SE. Note that even for inharmonic SFS shifts, the AC cue shared 83% of the variance with the average empirical responses, presumably highlighting the residual periodicity information comprised in shifts of inharmonic sounds.

To rule out the possibility that the success of the model to fit the behavioral data arose as a consequence of overfitting, we fitted the same model to shuffled data obtained by randomizing the indices of the shift size factor (using the same index randomization for all participants and weightings) for 1000 distinct index permutations. If our results were merely a consequence of overfitting through the optimization of cue weights, then the model should still successfully be fitted to human data for shuffled cue values. It turned out that the best (i.e., 99th percentiles) of the simulated correlations had *R*^2^ values that were at least 20 percentage points lower compared to the actual *R*^2^ values (see Supplementary Tab. 6). Hence, our model fitting procedure generally did not yield overfitting, but rather allowed us to capture pertinent aspects of the auditory cues underlying perceptual responses.

### Modeling inter-individual differences in cue weighting

The model computed the same three auditory cues for all listeners, but allowed for inter-individual differences in the weighting of these cues when fitting model predictions to behavioral responses. To qualitatively illustrate the inter-individual differences across listener-fitted model, Fig. 8 shows the optimal weights across participants and experimental conditions, as well as an estimate of the spread of these estimates obtained by a bootstrapping technique (see Methods). For harmonic SFS shifts, participants’ behavior was best described by the AC cue with very little inter-individual differences across participants. This is generally consistent with the consensus idea that comparing two harmonic complex tones is mediated by a periodicity analysis, the core mechanism of most current pitch models. There was one exception, namely a participant who more strongly relied on CCres cues, but with high uncertainty in the selection of weights, because almost all the scattered bootstrap samples in the plot stem from this one participant, see Supplementary Fig. 3. More spread across cues weights was obtained in the other conditions, which could be explored for the first time with the generalized Shepard tones. For inharmonic SFS shifts, AC and CCunres were weighted most strongly across the group of listeners, yielding considerable spread of weightings across participants. For harmonic SE shifts, listeners most strongly weighed the CCres cue, even though one participant mostly relied on the CCunres cue. For inharmonic sounds, the weights were generally wide-spread across participants, suggesting large inter-individual differences. For inharmonic SE shifts, the distribution of optimal weights was roughly centred in the middle of the triangular plot, which suggests that most participants tended to assign roughly equal weights all three types of cues.

**Fig 8.**
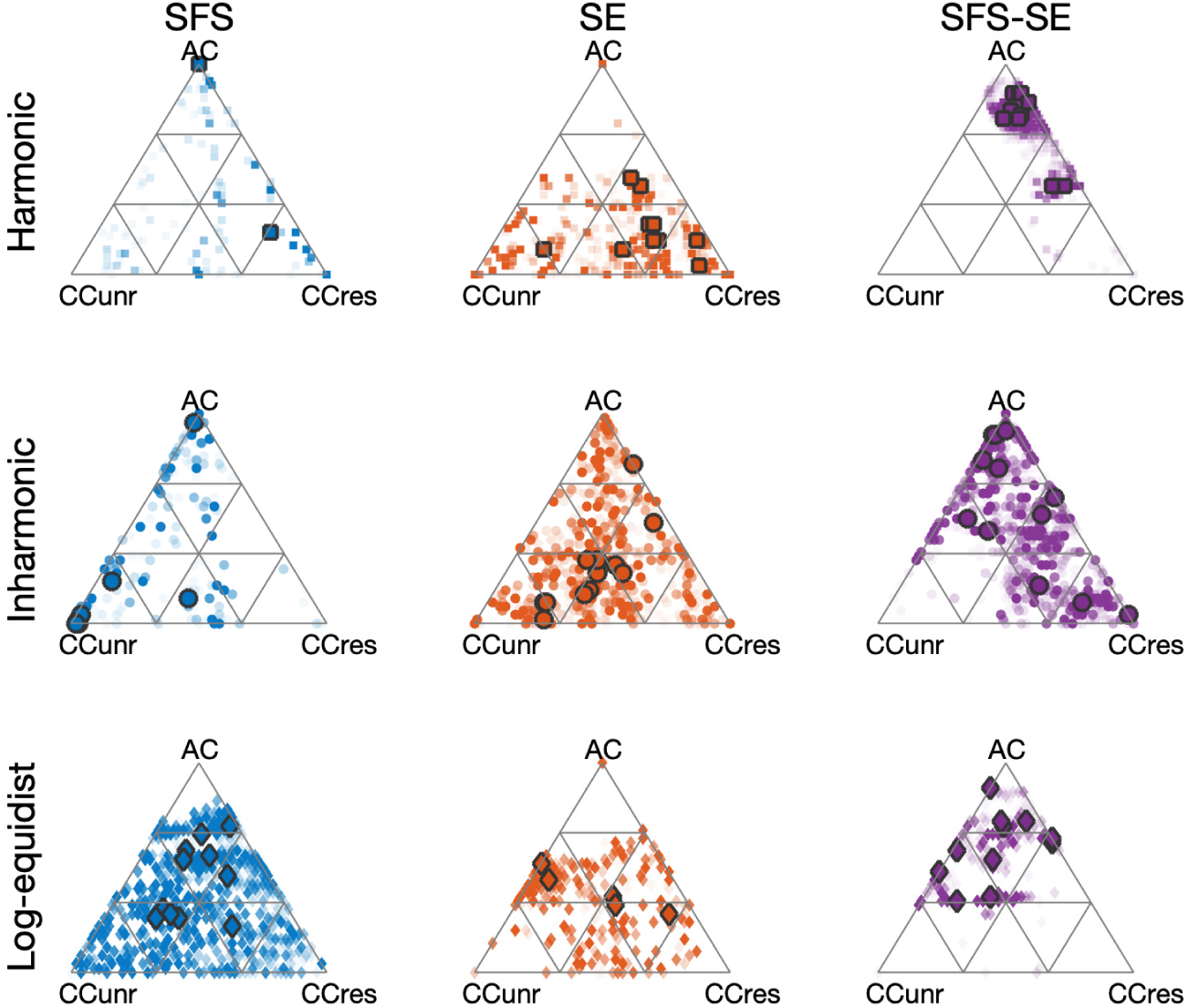
Cue weights for all conditions and participants with panels sorted as in Fig. 7. Constrained to sum up to one, cue weights live on a two-dimensional hyperplane with triangular borders. Points in the triangular space correspond to weightings of the three cues AC, CCres, and CCunres (weights sum up to one). Large symbols correspond to the medians of the optimal weighting as computed from 2000 bootstrap iterations. Transparent symbols highlight spread of bootstrap samples (the less transparent, the more overlap from different bootstrap iterations).

We now describe the two-dimensional SFS-SE shifts. These conditions are of special interest, because listeners may have implicitly relied on different weight combinations when faced with conflicting acoustic cues. This is qualitatively what was observed. In the harmonic SFS-SE case, two clusters of participants were observed along the AC-CCres continuum: most listeners predominantly relied on AC, the universal cue for harmonic SFS shift, while two participants also took into account CCres, the most weighed cue for harmonic SE shifts. A much more wide-spread clustering of participant was observed for the inharmonic SFS-SE case, although the CCunres cue was not predominantly used by any of the participants in this condition. Note that the inharmonic one-dimensional SFS or SE shifts were also the ones with the larger spread. Finally, for the log-equidistant SFS-SE case, AC and CCunres cues appeared to be dominant.

## Discussion

We introduced a novel auditory stimulus to probe the effect of different acoustic and auditory cues on perceptual judgments of the “low” to “high” scale, which is commonly associated to pitch perception. Our stimuli were generalized versions of classic Shepard tones. The original Shepard tones proved pivotal in trying to disentangle the contribution of two putative cues to pitch termed chroma and pitch height [49, 61]. Our generalized Shepard tones further allowed the independent manipulation of two types of acoustic dimensions, SFS and SE, which could be crossed with properties of harmonicity and inharmonicity. These acoustic dimensions were chosen based on the physics of natural sounds, with SFS related to source excitation and SE related to filter resonance [12]. By design, generalized Shepard tones displayed cyclicity over frequency shifts, the central feature of the original Shepard tones. We further demonstrated ambiguity for frequency shifts corresponding to a half frequency cycle. Moreover, because of the inherent symmetry of generalized Shepard tones, we could investigate in a parametrical manner the synergistic or antagonistic interactions of changes along the SFS and SE dimensions. A first finding was that, even in the case of conflict between SFS and SE, cues were implicitly combined for the perceptual reports. A detailed examination of the interactions between SFS, SE, harmonicity, and two types of inharmonicity demonstrated that the acoustic dimensions were not equally effective for all types of sounds. To quantify this observation, a computational model was designed, based on three estimated auditory cues. These auditory cues were chosen based on perceptual models of pitch (AC), frequency-shift detection (CCres), and timbre perception (CCunres). The model was highly effective at accounting for the behavioral data. It further specified the implicit weighting of cues in each experimental condition, revealing effects of sound type and inter-individual differences.

### What did participants judge in the behavioral task?

Our experimental design involved presenting two sounds, and asking listeners to report whether they heard a shift “up” or “down”. We explicitly stated in the instructions that the shift could be described as a change in pitch or in timbre brightness. A first issue is, then, what did listeners actually judge: pitch, or brightness? We will argue here and in rest of this Discussion that this distinction, while theoretically relevant, is not central to the interpretation of our experimental data and modeling results.

A large body of literature has now reported interactions between pitch and brightness in terms of discrimination experiments [8, 32, 35] and interval estimation [62, 63]. In spite of the labels provided to listeners in these experiments, it is intrinsically impossible to know whether listeners judged “pitch” or “brightness” to perform the tasks successfully. Moreover, such a sharp distinction becomes a moot point in light of the strong interactions observed.

A complementary approach consists of relating listeners’ behavior to auditory cues, which can be mathematically defined irrespective of the label attached to them and hypothetical underlying perceptual dimension. A long history of models have contributed to clarify and produce testable hypotheses about pitch (reviewed in [5]). It is precisely in this spirit that we designed our own computational model, which shares many features with existing ones. One novelty is that our model included several candidate auditory cues, all contributing to a single decision process on an equal footing, but with the weight of each cue left free to vary depending on the stimulus type and listener. Experimentally, this approach proved successful to account for a large behavioral dataset, further suggesting that a combination of cues was necessary and sufficient to account for the perceptual “up” vs. “down” judgments provided by listeners.

Did participants access a perceptual correlate of each cue, and combine them at the decisional level, or was there rather a unitary perceptual dimension underlying their response? A notable feature of our findings concerns the cases where acoustic dimensions, and thus presumably auditory cues, were put in conflict. In these experiments (Exp. 2 and Exp. 3B), listeners had four response choices: “up”; “both (up dominant)”; “both (down dominant)”, “down”. Thus, they were encouraged to report when they heard two distinct cues signaling two different shift directions. However, “both” responses were only rarely reported, which was not a priori obvious. For instance, Singh and Hirsh [33] provided labels of pitch and timbre changes to report judgments of concurrent SFS and SE shifts, and listeners appeared to be able to use such labels independently. Interestingly, here “both” responses were not dominant even when responses were split between around 50% up and 50% down reports, suggesting a fundamental ambiguity in the stimulus similar to the tritone paradox for the original Shepard tones [49, 51–53]. Unlike the tritone paradox case, here the ambiguity was caused by opposing changes along acoustic dimensions, and not only by ambiguous cues in either domain. Such an ambiguity did not seem to clearly reach metacognitive awareness, as “both” responses were rare even in ambiguous conditions. This finding parallels recent results using single-cue ambiguity with the original Shepard tones [64]. Moreover, a recent imaging study [61] also used Shepard tones with conflicting pitch chroma and pitch height shifts, akin to SFS and SE shifts. They also reported unitary perceptual judgements. In their study, perceptual judgements were overwhelmingly dominated by pitch chroma. But, as pointed out by the authors, this was likely due to a lack of parametric flexibility of the original Shepard tones. Our generalized paradigm extends these results, including conditions similar to those of [61]. Consistent with their results, AC cues dominated harmonic SFS shifts. Yet, other conditions such as inharmonic SE shifts were strongly dominated by other cues derived from the SE shifts.

In summary, the present results taken together with the well-established interaction between acoustic cues suggest that changes along the frequency axis, whatever their acoustic nature, may be projected onto a “low” to “high” perceptual dimension when the task is to track their changes over time. For clarity, we will now describe our results in terms of frequency change perception, and not pitch or brightness. Note that this terminology is not meant to imply that listeners did not in fact perform “pitch” judgments. Rather, it is intended to distinguish the remaining discussion points from this first, basic issue.

### The role of acoustic cues in frequency change perception

Physical objects produce sounds that evolve over time, either because a vibratory source changes its behavior or because resonant bodies excited by the source change their shape. Such ecologically-relevant events lead to changes of SFS and SE properties. The present behavioral results show that both types of changes impact the perception of frequency change, but to a different extent depending on the nature of the sound. Exp. 1 tested one-dimensional SFS and SE cues for harmonic and inharmonic series of frequency components. Harmonicity was found to mediate the relative strength of the perceived shift. For harmonic sounds, changes over the SFS dimension yielded steeper slopes as a function of shift size compared to changes over the SE dimension. For inharmonic sounds, the reverse was true, with changes along the SE dimension producing steeper slopes. Exp. 2 tested two-dimensional changes on both the SFS and SE dimensions. Here again, harmonicity mediated the results, in a manner consistent with Exp. 1. The SE dimension had a much smaller effects on shift perception for harmonic sounds compared to inharmonic sounds. Exp. 3, using a different type of sounds, replicated the dominant effects of SE changes for inharmonic sounds. In summary, these findings suggest that the perception of frequency changes was dominated by SFS cues for harmonic sounds, but not necessarily for inharmonic sounds.

Why would different acoustic cues be used for frequency-change tracking depending on the harmonicity of the sounds? Speculatively, it can be remarked that harmonicity plays an important role in auditory scene analysis [65–67]. In particular, harmonicity favors the binding of frequency components within and across sounds [26, 65]. Local frequency changes in SFS may then be more informative for harmonic sounds, whereas coarser SE cues may take over for inharmonic changes. However, a recent study tested pitch discrimination with interference of envelope variation and did not observe a mediating role of harmonicity in all conditions [26]. Instead, they only found a mediating role for very large intervals (9 st) or in the case when there was no overlap between the frequency components of the two sounds, but not for natural speech or musical instrument sounds. Future work should attempt to reconcile these apparently divergent findings on the role of harmonicity in acoustic cue weighting.

### The role of auditory cues in frequency change perception

To bridge the gap between the acoustic description of stimuli and the behavioral judgments by listeners, we used a computational model estimating a variety of auditory cues available to auditory processing. Mostly, this involved simulating the effects of peripheral filtering on frequency encoding [59], and extracting three types of cues inspired by models of pitch perception (AC, [5]), frequency-shift detection (CCres, [27]), or timbre perception (CCunres).

The choice of cues was somewhat arbitrary. It was nevertheless partially validated by two experimental observations. First, the model achieved a high accuracy in predicting behavioral data, with more than 90% of the variance explained in most cases. The three cues selected were thus sufficient to predict frequency change perception with generalized Shepard tones. The three cues were also only weakly correlated with each other, showing that they were not redundant. The only weak but significant correlation that was observed was between CCres and CCunres cues. These cues shared the same computational principle, cross-correlation between successive excitation patterns, but operated on different part of the frequency scale. CCres was focused on resolved harmonic while CCunres was focused on unresolved harmonics. The distinction between resolved and unresolved harmonics may not, in fact, be clear cut [68]. Thus, we are using the CCres and CCunres cues a a proxy for processes that capture distinct aspects of the excitation pattern, without necessarily being completely independent. Finally, all cues contributed in various combinations across experimental conditions and listeners, suggesting they were all necessary.

The auditory cues estimated could be mapped to some extent to the acoustic SFS and SE dimensions. For harmonic sounds, the AC cues provided a reliable estimation of sound periodicity and hence captured the specific pattern of frequency components for harmonic SFS. This may explain why AC dominated perceptual judgments for harmonic SFS shifts. Given the duality of spectral and temporal fine-structure information, we suspect that the AC cue could also be implemented based on SFS information alone [69], but not with the common Gammatone front end we chose to use for all cues estimated. For the CCres cue, because of the cross-correlation operation, we argue that this auditory cue is reflecting local shifts in the SFS rather than global pitch estimates, but this remains speculative. We acknowledge that the information derived from resolved harmonics is dominant for pitch perception as well [4, 69, 70]. The CCunres cue, finally, is most clearly linked to SE.

### Inter-individual in frequency change perception

Another novel finding revealed by the multi-cue model concerns inter-individual differences. To simulate a perceptual decision, our model assigned weights to each auditory cue before combining them through simple summation (normalized by each cue’s variance). Weights were individually fitted to each listener. Results showed that weights indeed varied across listeners, especially for inharmonic sounds and for cases with conflicting SFS and SE cues. The differences observed were not due to model overfitting, but rather represented true inter-individual differences in the weighting of biologically-plausible auditory cues when faced with a task of frequency change perception.

Intriguingly, there are already several lines of evidence demonstrating large interindividual differences in pitch perception. To start with, the perception of the tritone paradox with Shepard tones famously shows differences in bias across listeners (which *f*_0_ is perceived as higher). These inter-individual differences have been claimed to reflect the auditory experience of each listener, and in particular their linguistic background [71] although this has been contested [72, 73]. Experiments with simultaneous changes in SE and SFS often emphasize a large inter-individual variability in the results, even when independence between dimensions or labels were observed [30, 33]. In an experimental setting closer to our own, several studies have introduced conflicts between what has been termed “low-pitch” or “global pitch” on the one hand, and “spectral pitch” on the other hand [22, 74, 75]. In these studies, frequency components were changed across successive tones so that the missing fundamental changed in one direction whereas the frequency of each component changed in the opposite direction. In our terminology, this would approximately correspond to a conflict between SFS and SE cues. For our model, this would be a conflict between AC and CCres cues. Substantial interindividual differences were found in such tasks [75], with listeners described as holistic (AC cues dominant) or analytic (CCres cues dominant). These differences were again ascribed to auditory experience, in this case to musicianship [75, 76]). In our results, pronounced individual differences between listeners favoring different weightings of AC and CCres cues were observed in particular for the inharmonic two-dimensional SFS and SE shifts. This could suggest that the holistic and analytic distinction demonstrated by previous studies might be even more pertinent for the case of inharmonic sounds.

### Pitch as an adaptive combinations of sensory cues

Based on the experimental data and modeling results, we argue that frequency changes are tracked thanks to a perceptual dimension derived from the combination of different auditory cues. At least parts of the acoustic cue manipulations that were investigated would conventionally be described as changes in pitch. Thus, as we observed that all judgements involving generalized Shepard tones mapped to a single perceptual “low” to “high” dimension, then this dimension may perhaps be described as a generalization of pitch. A nuance may be introduced between such a dimension and musical pitch, which for Western music has the added constraints of being able to support interval perception, melodic patterns discrimination, and be robust to frequency transposition, none of which were tested here. But, as reviewed in the Introduction, the distinction is not so clear in all musical idioms, and timbre has been shown to share the ability to convey musical melodies [43]. In summary, a broader and more speculative claim of our findings is that pitch is a perceptual dimension useful to track frequency changes, which may be seamlessly constructed from a variety of acoustic cues as relevant to each sound class and listener.

Such a claim is clearly somewhat unorthodox with respect to the long and respected tradition of investigations about pitch. However, multi-stage pitch models have already been proposed to account of temporal integration properties [77]. Moreover, it can be pointed out that a multi-cue model would put pitch on par with another fundamental dimension of auditory perception, namely spatial location. The perception of subjective spatial location tracks the physical spatial position of objects in the world. However, spatial position has no direct correlate in the sensory receptor for hearing, the cochlea, which is tonotopically organized in terms of frequency channels rather than space. As a consequence, location needs to be perceptually constructed from a combination of binaural and monaural cues that are very different in nature from each other. Binaural cues compare the sensory information between the two ears, looking for discrepancies introduced by differences in acoustic paths from the object to the ears, whereas monaural cues are based on a location-based spectral filtering of the acoustic signal. Within binaural cues, interaural differences in timing (ITDs) and in level (ILDs) are processed in separate parts of the auditory pathways. Presented monaurally, the level differences corresponding to ecological ILDs are clearly heard as differences in loudness. However, ILD and ITDs are seamlessly combined into the perception of spatial location. These general features of auditory spatial location perception are qualitatively equivalent to our claim of a “unitary” pitch perception mechanism based on multiple cues.

The parallel between our results and the perception of spatial location can even be made more specific. In the implicit combination of binaural cues to derive location, the weighing of ITDs and ILDs is stimulus-specific: to localize low-frequency noise bands, ITD will be weighed more than ILDs, while the converse is true for high-frequency noise bands [78]. Thus, cue weighting depends on the stimulus. Conflict between ITD and ILD cues has also been investigated, and it has been observed that they can be precisely counterbalanced so that an ITD pointing e.g. to the left combined with an ILD pointing to the right results in a sound perceived in the middle [78]. Thus, they contribute to a unitary dimension. Finally, for broadband noise, there are distinct inter-individual differences in ITD and ILD weighting [79]. Similar effects of cue integration have been found for the lateralization of sounds with incongruent ITDs of signal temporal fine structure and temporal envelope [80], two different acoustic cues which are processed independently in the early stages of the auditory system, including pronounced inter-individual differences [81]. Monaural cues are also known to be strongly idiosyncratic, to the point that monaural cues from someone else’s ear are heard as timbre changes, but become transparent cues to location for one’s own ears. Moreover, it appears that they can be learnt through experience [82]. Thus, auditory cues to spatial location are derived and combined in a listener-specific manner.

### Conclusion

The present behavioral data and computational results suggest that, just as is the case for spatial location, a unitary perceptual dimension of “pitch” is derived from a set of various acoustic cues in a sound-specific and listener-specific manner. There are no direct correlates of the physical properties of sound-producing objects on the cochlea, but there are many indirect correlates related to the frequency content of sounds. A perceptual dimension may then be constructed from several cues, with a stimulus-specific weighting of cues and additional inter-individual differences in the weighting. It might be tempting to think that pitch is a gift from above so that we can enjoy Bach chorales, or that it neatly maps to a single acoustic cue, but this is probably not how evolution works. Instead, we speculate that a compound perceptual dimension tracking frequency-changes has adaptive value for hearing in complex and ever-changing environments.

## Methods

### Participants

Participants were recruited via ads at the online job board of the University of Oldenburg and received monetary compensation for their time. All experiments tested participants of self-reported normal hearing. In Exp. 1, thirteen participants were tested, but one participant failed to meet the screening test (see Procedure) and was not considered in the analysis; the remaining twelve participants had a median age of 25 years (range: 20–56 years). Considering musical training of participants (which we did not specifically attempt to balance), there were two participants without any musical training; the ten participants with musical training were amateur musicians with a median of 11 years of training on their main instrument (range: 5–19 years) and a median of 54 points (range: 27–73) on the Goldsmiths-Musical Sophistication Index (Gold-MSI) musical training subscale [83].

In Exp. 2, twelve participants were tested and all passed the screening test. Participants had a median age of 24.5 years (range: 21–30 years). There were three participants without any musical training; the remaining participants were amateur musicians with a median of 10 years of training on their main instrument (range: 6–20 years) and a median of 54 points (range: 30–82) on the Gold-MSI musical training subscale.

In Exp. 3, twelve participants were tested (all passed the screening). Participants had a median age of 25 years (range: 19–29 years). There were four participants without any musical training; the remaining participants were amateur musicians with a median of 10 years of training on their main instrument (range: 3–15 years) and a median of 43 points (range: 30–62) on the Gold-MSI musical training subscale.

### Stimuli

The experiment presented artificially synthesized tone complexes that were created using additive synthesis implemented in MATLAB (www.mathworks.com). Tone complexes were generated with fundamental frequency *f*_0_ randomly drawn between 64 and 128 Hz and partials covered the range up to 16 kHz. In the harmonic condition, partial tones were at integer ratios *f_k_* = *f*_0_ · *k* Hz (i.e., equidistant on a linear scale). In the inharmonic condition, partial tone frequencies were jittered such that 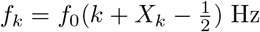, where for all *k, X_k_* is a uniformly distributed random variable with values between zero and one, see [25]. The same instantiation of X_k_ was used for any pair of sounds within trials of the inharmonic condition.

To create an SFS scale that was circular with regards to pitch chroma, the amplitudes of odd harmonics were attenuated as a function of the scale position in the octave, see Fig. 1. At scale position 0, odd harmonics were attenuated by 22 dB and raised by 2 dB for every semitone (st) step. This attenuation is smaller than the value of 3.5 dB in [55], but in combination with SE manipulation appeared to yield favourable results. Hence, after 11 st steps, odd and even harmonics were of equal amplitude.

The SFS manipulation was combined with an SE manipulation that has been shown to yield circular brightness perception [45]. A global spectral envelope was defined as Gaussian distribution as a function of log-frequency with a mean three octaves above f0 and a standard deviation of one octave. This global envelope was multiplied with a more fine-grained spectrally periodic envelope consisting of superpositions of octave-shifted Gaussians with a more narrow standard deviation of 0.15 (and 0.2 in Exp. 3 to account for the log-equidistant partials). The resulting spectral envelope hence was the product of superimposing a global and a local spectral envelope. Figure 1B shows an illustration. Sounds were synthesized with a duration of 125 ms including a 5 ms fade-in and fade-out using cosine ramps. Inter-trial sequences were used to minimize carry-over effects [45, 53]. These consisted of three sounds with a duration of 125 ms (including 5 ms raised cosine fades), separated by 125ms inter-stimulus-intervals and sounds had random spectra with 30 partial tones of random initial phase and random frequencies drawn uniformly from a linear scale between 50 Hz and 10 kHz. These tone complexes were synthesized with the same global spectral envelope that was used for the target tones in the subsequent trial.

In Exp. 1, every trial presented a pair of sounds, which was generated through unidimensional shifts of either SFS or SE . For the target attribute, the starting position on the circular SFS or SE scales was chosen at random integer scale positions from 0-11 st. The attribute that was kept fixed was held constant at shift position 0. In Exp. 2, stimuli consisted of two-dimensional shifts, that is, both SFS and SE were shifted simultaneously. Here, the starting positions of both the SFS and SE attributes were chosen at random.

Stimuli in Exp. 3A and B used equidistant partial tones on a logarithmic scale with a distance of four semitones, that is, three partials per octave. In Exp. 3A, unidimensional shifts were presented, similar to Exp. 1. In Exp. 3B, bidimensional shifts were presented, similar to Exp. 2. Sound examples can be found under https://uol.de/en/music-perception/sound-examples/generalized-shepard-tones-1/conflicting-spectral-cues

### Apparatus

Listeners were seated individually in a sound-proof lab and provided responses on a computer keyboard. Sounds were synthesized in MATLAB and were presented with an RME FIREFACE audio interface at an audio sampling frequency of 44.1 kHz and 24 bit resolution. Stimuli were presented diotically over Sennheiser I 200 headphones at 65 dBA sound pressure level, as calibrated by a Norsonic Nor140 sound-level meter with a G.R.A.S. IEC 60711 artificial ear to which the headphones were coupled.

### Procedure and design

The research reported in this manuscript was carried out according to the principles expressed in the Declaration of Helsinki and was approved by the ethics board of the University of Oldenburg. All experiments started with a screening task [45], comprising 40 SE trials with 2 st and 3 st shifts. The screening task was used to ensure that participants understood the task. Participants who did not achieve scores of more than 80% correct responses did not take part in the main experiment (which turned out to be the case only for one participant of Exp. 1).

The experimental instructions (originally formulated in German) explained that the task was to judge whether the second sound was perceived as “*higher or brighter (that is, going upwards) or lower or darker (that is, going downwards)*” in comparison to the first sound. For the two-dimensional shifts of Exps. 2 and 3B, instructions noted that it is possible that both directions can be heard at once, in which case participants were asked to indicate the dominant direction by selecting one of the four response alternatives: “down”, “both (down dominant)”, “both (up dominant)”, and “up”.

In Exp. 1, only one-dimensional shifts were presented. The experiment contained the factors of shift size (1-11 st), shift type (SFS, SE), and harmonicity (harmonic, inharmonic). There were 24 trials per condition with randomized scale positions of sound 1 that were split half across two sessions on separate days. Overall, this yielded 24×11×2×2 = 1056 trials per participant. Trials were presented in four blocks, with each block corresponding to one of the four shift type x harmonicity conditions. The presentation order of blocks was randomized and the presentation of blocks was separated by short breaks of around 5 minutes. Levels of the shift-size factor were presented in fully randomized order.

Experiment 2 contained the factors SFS shift size (1, 6, 11 st), SE shift size (1, 6, 11 st), and harmonicity (harmonic, inharmonic). There were 36 trials per conditions with randomized scale positions of sound 1. Overall, this yielded 36×3×3×2 = 648 trials in total. The harmonicity factor was presented blockwise with short breaks separating the blocks and the data were collected in one session. Participants could take in between trials, if requested.

In Exp. 3A, participants completed the unidimensional shifts, with shifts of the target attributes (SFS, SE) presented blockwise. The experiment contained the factors shift type (SFS, SE) and shift size (1,2, 3 for SFS; 1, 6, 11 for SE). There were 24 trials per condition with randomized scale positions of sound 1, overall yielding 24×2×3 = 144 trials. The shift type factor was presented blockwise with order of presentation randomized across participants. After a short break, participants went on to complete Exp. 3B, which contained the two factors SFS shift size (1, 2, 3 st) and SE shift size (1, 6, 11 st). As in Exp. 2, there were 36 trials per condition with randomized scale positions of sound 1, overall yielding 36×3×3 = 324 trials, which were presented in fully randomized order.

### Data analysis

Data were analysed in MATLAB (https://www.mathworks.com). In Exps. 2 and 3B, which comprised two-dimensional shifts, “both (up dominant)” responses were collapsed with “up” responses and “both (down dominant)” with “down” responses. This approach is justified by the generally low number of both responses, as visible in Supplementary Fig. 1.

Trial-level down responses were analyzed using generalized linear mixed-effect (GLME) models as implemented in the fitglme class, using a logit link function and a binomial distribution of the response variable. All models comprised by-participant and by-item random intercepts (the latter encoded the randomly chosen position of sound 1 along the circular SFS scale (see Fig. 1A). In Exp. 1, the shift factor was numerically coded without normalizing predictors. In Exp. 2, the SFS shift factor was coded numerically (without normalization) whereas for the SE factor, effects coding was used (corresponding to Fig. 3). In Exp. 3A, both shift factors were numerically coded (using factor levels 1, 2, and 3). In Exp. 3B, factors were coded analogously to Exp. 2 (see Fig. 4B), that is, the SFS factor was coded numerically and for the SE factor effects coding was used. For all other categorical predictors effects coding was used. The full results of the statistical models are provided in Tables 1-4 in the Supplementary Materials. The fitted statistical models are plotted in all results figures as solid lines. Square brackets in the text indicate 95% confidence (i.e., compatibility) intervals.

### Computation of acoustical cues

The two signals presented on every trial were processed by a 128-channel Gammatone filterbank spanning 20 to 16000 Hz. For the autocorrelation (AC) cue, a standard pitch estimator was used (de Cheveigné, 2005). Specifically, the AC functions were computed for every band of signals 1 and 2 and summed across bands. The maximal values of the resulting summed AC functions were used as *f*_0_ estimates for signals 1 and 2, see Fig. 5. The *f*_0_ search range was constrained between 64 and 128 Hz, which is the *f*_0_ range used in Exps. 1 and 2. Formally, the AC cue hence can be written as

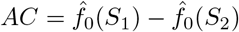

with 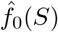 denoting the *f*_0_ estimate of signal *S*.

The cross-correlation (CC) cues extracted shifts between the excitation patterns of signals 1 and 2. The CCres and CCunres cues were computed identically, but considered the presumed resolved and unresolved portions of the excitation pattern, respectively, with the cutoff point chosen at the geometric mean of the 10th harmonic [84] of sounds in Exps 1-2 (905 Hz). Note that other implementations of the cutoff frequency, chosen as the exact frequency of the tenth partial of the sound with lower *f*_0_ on every trial or even chosen randomly in the range between 640 and 1280 Hz yielded qualitatively and quantitatively very similar results.

The root-mean-square sound levels were computed for every of the 128 channels of the filterbank, and the overall profile was peak-normalized at 0 dB across channels and thresholded at −30 dB (i.e., levels below −30 dB were set to −30 dB). The cross-correlation function between the two resulting excitation patters of signals 1 and 2 was computed. Subsequently, the first moment of the CC function was computed in the heuristically chosen range of −5 to +5 ERBs, corresponding to a centroid of the CC function in ERB-lag units.

Formally, the CC cue can thus be written as

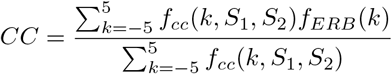

with *f_cc_*(., *S*_1_, *S*_2_) denoting the cross-correlation function of the excitation patterns of sounds *S*_1_ and *S*_2_ on a single trial. We did not use the maximal value of the CC function (which would have been analogous to how the AC function was evaluated), because for the present stimuli with a fixed global spectral envelope in every trial, the maximal value turned out to constantly occur at a lag of zero, calling for a more sensitive measure. Instead, we used the CC centroid as a continuous estimate of the strength of downward vs. upward frequency movement with negative CC centroids indicating dominant downward movement and positive CC centroids indicating upward movement.

### Cue weighting

Similar to classic maximum likelihood approaches of multisensory integration [85], our model integrated the sum of individual estimates corrected by their variance. To capture the potentially stimulus- and participant-dependent weighting of AC, CCres and CCunres cues [75], three weighting parameters *α*_1/2/3_ allowed to assess which type of cues would best correspond with participants’ behavior. Let *σ*^2^ denote the variance of the estimators specific to the stimuli computed individually for every of the three experiments. Then the three cues were integrated using the weighting factors *α*_1_ + *α*_2_ + *α*_3_ = 1 with the following response criterion (“down” response, if valid):

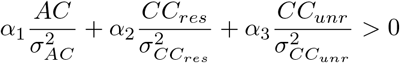

For every participant and experimental block, we computed the optimal combination of weights, yielding the largest Pearson correlation between individual response pattern (proportion “down” responses) and corresponding model responses by using a grid-search with 2% resolution (0 ≤ *α*_1/2/3_ ≤ 1 such that *α*_1_ + *α*_2_ + *α*_3_ = 1), yielding 1326 different parameter triples. Because the set of non-negative weights live on a hyperplane, they can be represented as part of a triangle in two dimensions (Fig. 6B-C).

The stability of the derived weights was assessed using bootstrapping with 2000 iterations. In every iteration, the response of individual participants was perturbed by only computing response patterns from a random subset of trials from the respective experiment (same number of trials as in the original experiment but drawn with replacement). The procedure for selecting weights described above was subsequently conducted for every bootstrap iteration (each containing a different perturbation of participant’s responses). Fig. 8 presents the median of the derived weights (large symbols) across all iterations in addition to all 2000 derived weights plotted with transparent symbols. That is, the less transparent a region in the figure is, the more often weights from different bootstrap iterations overlap in that region.

To measure how well the fitted model also accounted for the global pattern of responses (data averaged across participants), we averaged the models fitted to individual human participants. The resulting response patterns is shown in Fig. 7 with *R*^2^ values provided in the figure. For Exp. 3a, there are only three data points, thus correlation coefficients were not considered reliable and omitted in the figure.

## Acknowledgements

KS is supported by a Freigeist Fellowship of the Volkswagen Foundation.

The authors thank Alain de Cheveigné and Laurent Demany for valuable comments on the manuscript.

## Author contributions statement

KS, JG, and DP designed the experiments. KS collected the experimental data and conducted the data analysis and computational modeling. All authors discussed experimental and modeling results. KS and DP devised the first full draft of the manuscript. All authors discussed and approved the final version of the manuscript.

