## Supplementary Materials for "A unitary model of auditory frequency change perception"

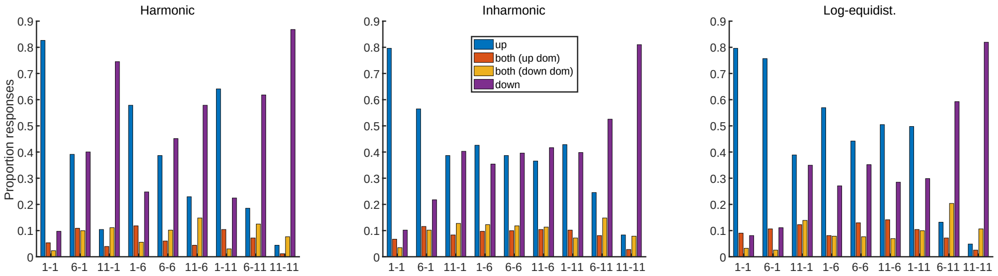


**Fig 1.** Average proportion of responses for sounds with harmonic (left), inharmonic (middle), and log-equidistant partial series (right) for every of the four response categories. Description of the x-axis corresponds to the size of the SFS and SE shift in semitones.

### GLME models

**Table 1.** Exp. 1. Results from the GLME model with the independent variables harmonicity ("harm", harmonic vs. inharmonic), shift type ("type", SFS vs. EN), and shift size ("size", 1, 2,..., 11), as well as interaction terms (indicated by ":"). Model statistics: AIC = 62648, $R^2 = .31$.

| Variable | $\beta$ | CI low | CI high | t-value | p-value |
|---|---|---|---|---|---|
| intercept | -2.02 | -2.32 | -1.71 | -13.04 | ¡.001 |
| harm | -0.26 | -0.35 | -0.17 | -5.73 | ¡.001 |
| type | 0.01 | -0.08 | 0.1 | 0.17 | 0.865 |
| size | 0.31 | 0.3 | 0.32 | 45.23 | ¡.001 |
| harm:type | -0.52 | -0.61 | -0.43 | -11.39 | ¡.001 |
| harm:size | 0.06 | 0.04 | 0.07 | 8.49 | ¡.001 |
| type:size | 0 | -0.01 | 0.01 | -0.07 | 0.944 |
| harm:type:size | 0.1 | 0.08 | 0.11 | 14.12 | ¡.001 |

**Table 2.** Exp. 2. Results from the GLME model with the independent variables harmonicity ("harm", harmonic vs. inharmonic), SFS shift size ("FS", 1,6,11), first SE shift dummy variable ("SE1", 1 vs. 11), second SE shift dummy variable ("SE6", 6 vs. 11), as well as interaction terms (indicated by ":"). Model statistics: AIC = 36416, $R^2 = .45$.

| Variable | $\beta$ | CI low | CI high | t-value | p-value |
|---|---|---|---|---|---|
| intercept | -1.28 | -1.63 | -0.93 | -7.17 | ¡.001 |
| harm | -0.39 | -0.49 | -0.3 | -8 | ¡.001 |
| SFS | 0.25 | 0.23 | 0.26 | 32.74 | ¡.001 |
| SE1 | -0.99 | -1.14 | -0.85 | -13.2 | ¡.001 |
| SE6 | 0.7 | 0.57 | 0.83 | 10.66 | ¡.001 |
| harm:FS | 0.09 | 0.08 | 0.11 | 12.81 | ¡.001 |
| harm:SE1 | 0.22 | 0.07 | 0.37 | 2.9 | 0.004 |
| harm:SE6 | -0.06 | -0.19 | 0.06 | -0.99 | 0.322 |
| FS:SE1 | 0.06 | 0.03 | 0.08 | 5.23 | ¡.001 |
| FS:SE6 | -0.14 | -0.16 | -0.12 | -14.33 | ¡.001 |
| harm:FS:SE1 | 0 | -0.02 | 0.02 | 0.3 | 0.765 |
| harm:FS:SE6 | -0.01 | -0.03 | 0.01 | -0.95 | 0.341 |

**Table 3.** Exp. 3A. Results from the GLME model with the independent variables shift type ("type", SFS vs. EN), shift size ("size", 1,2,3 for SFS and 1,6,11 for EN), and interaction terms (indicated by ":"). Model statistics: AIC = 7946.9, $R^2 = .35$.

| Variable | $\beta$ | CI low | CI high | t-value | p-value |
|---|---|---|---|---|---|
| intercept | -2.55 | -3.01 | -2.1 | -10.98 | ¡.001 |
| type | -0.92 | -1.24 | -0.61 | -5.79 | ¡.001 |
| size | 1.26 | 1.11 | 1.41 | 16.5 | ¡.001 |
| type:size | 0.52 | 0.37 | 0.67 | 6.95 | ¡.001 |

**Table 4.** Exp. 3B. Results from the GLME model with the independent variables FS shift size ("FS", 1, 2, 3), first SE shift dummy variable ("SE1", 1 vs. 11), second SE shift dummy variable ("SE6", 6 vs. 11), as well as interaction terms (indicated by ":"). Model statistics: AIC = 19243, $R^2 = .63$.

| Variable | $\beta$ | CI low | CI high | t-value | p-value |
|---|---|---|---|---|---|
| intercept | -2.25 | -2.81 | -1.68 | -7.78 | ¡.001 |
| SFS | 1.01 | 0.9 | 1.12 | 17.76 | ¡.001 |
| SE1 | -1.76 | -2.13 | -1.4 | -9.45 | ¡.001 |
| SE6 | 1.65 | 1.36 | 1.95 | 10.83 | ¡.001 |
| FS:SE1 | 0.25 | 0.09 | 0.41 | 3.1 | 0.002 |
| FS:SE6 | -1 | -1.14 | -0.86 | -13.9 | ¡.001 |

### Computational modeling

**Table 5.** $R^2$ values from Pearson correlation coefficients between raw feature values and average human listeners for AC, CCres, and CCunr features as depicted in Fig. 1. Values in bold font show significant entries according to Bonferroni correction $(0.05/9 = 0.0056.)$

| | | SFS | SE | SFS-SE |
|---|---|---|---|---|
| AC | Harmonic | **.97** | .01 | **.88** |
| | Inharmonic | **.83** | .00 | **.75** |
| | Log-equidist | .93 | .99 | .30 |
| CCres | Harmonic | **.75** | **.96** | .58 |
| | Inharmonic | **.83** | **.87** | **.93** |
| | Log-equidist | .61 | .99 | .06 |
| CCunres | Harmonic | .00 | **.91** | .06 |
| | Inharmonic | .39 | **.89** | .52 |
| | Log-equidist | 1.0 | .88 | .66 |

**Table 6.** $R^2$ values for 99th percentile of Pearson correlation coefficients between empirical data and model fitted with randomly permuted shift factor indices, measured over 1000 random permutations. Note that there are $9! = 362.880$ permutations of the set $\{1, ..., 9\}$ and $11! = 39.916.800$ permutations of $\{1, ..., 11\}$, hence it is highly unlikely that the non-permuted index set is part of the simulation.

| | SFS | SE | SFS-SE |
|---|---|---|---|
| Harmonic | .50 | 0.42 | .61 |
| Inharmonic | .40 | .44 | .67 |
| Log-equidist | .97 | .95 | .56 |

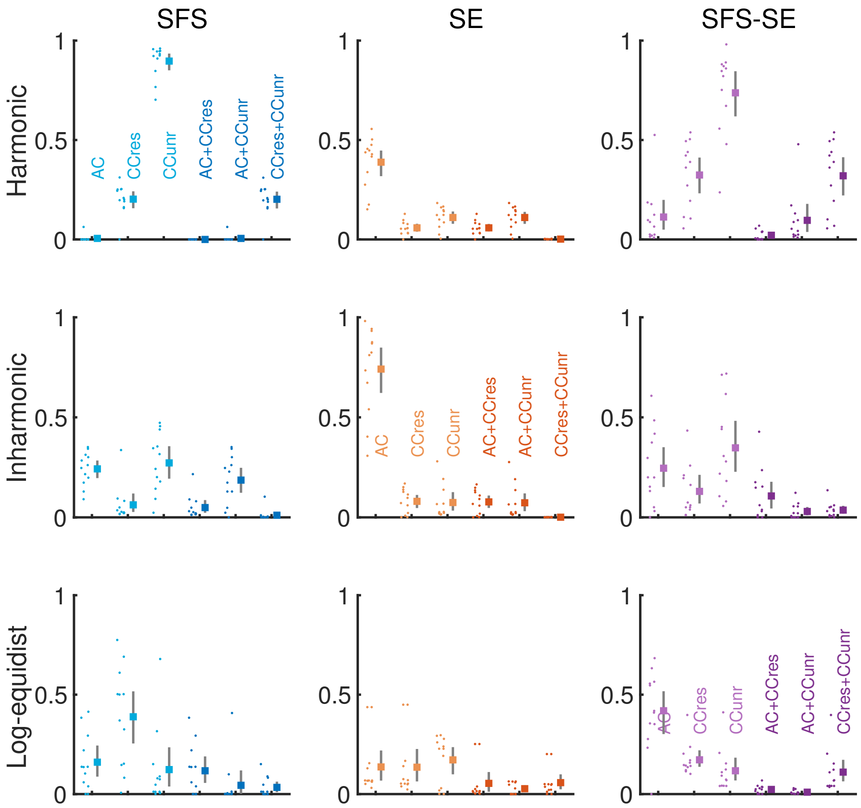


**Fig 2.** Distribution of paired differences of R2 values (displayed on the y-axis) between three-dimensional complete model (AC+CCres+CCunres) and lower-dimensional incomplete model variants (x-axis, see panels on the diagonal for labels). Panels sorted according to fine structure type (rows) and acoustic shift dimensions (columns). Dots correspond to differences for individual participants, square symbols to mean; error bars indicate boostrapped 95% confidence intervals. Of particular interest are the two-dimensional shifts displayed on the rightmost side. Here, it is visible that for the harmonic case, the fit of the AC+CCres model is indistinguishable from the full model. For the inharmonic case, however, the AC+CCres variant exhibits poorer fit (CIs non-overlapping with zero). That is, the full model is necessary to account for the general case of harmonic and inharmonic sounds.

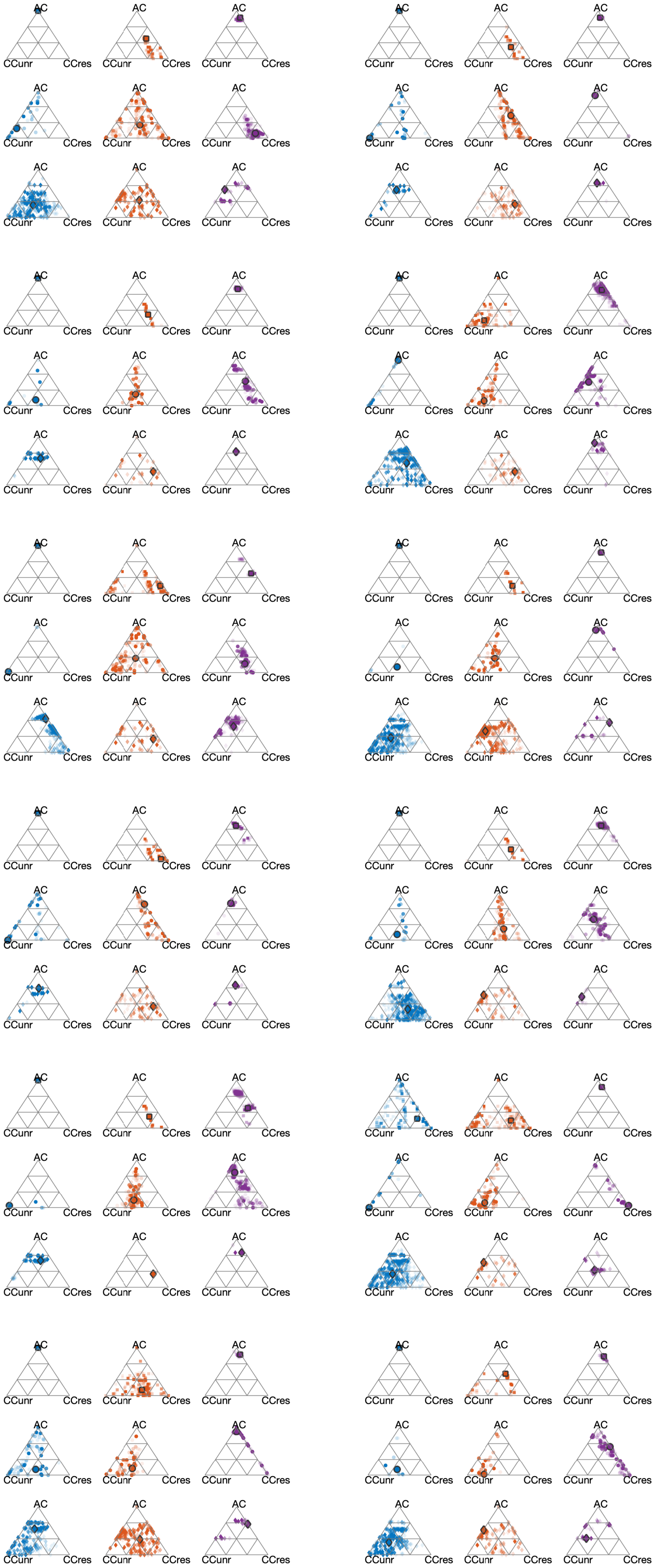

**Fig 3.** Boostrapped cue weightings for all individual participants.
